# Dynamic organizational strategies of multidomain glycosyltransferases revealed by high-speed AFM and solution biophysics

**DOI:** 10.64898/2026.06.04.729992

**Authors:** Hirokazu Yagi, You-Rong Lin, Yuki Kanaoka, Fumiko Umezawa, Akemi Kim, Kotaro Tomuro, Ken Morishima, Atsuji Kodama, Kentaro Ishii, Susumu Uchiyama, Tadashi Satoh, Masaaki Sugiyama, Takayuki Uchihashi, Koichi Kato

## Abstract

Glycosyltransferases often contain multiple structural modules that contribute to substrate recognition, catalytic coordination, and higher-order molecular organization. However, how multidomain glycosyltransferases dynamically organize their catalytic domains in solution remains poorly understood. Here, we investigated the assembly states and conformational dynamics of POMGNT2, LARGE1, K4CP, and L137 using high-speed atomic force microscopy (HS-AFM) integrated with complementary solution biophysical analyses.

POMGNT2 formed a stable dimeric architecture with limited large-scale conformational fluctuation, consistent with its role in site-selective substrate recognition. In contrast, LARGE1 and K4CP exhibited concentration-dependent and heterogeneous assembly behavior. K4CP displayed pronounced open–closed interdomain motion and substrate-dependent conformational compaction, indicating dynamic catalytic-domain reorganization during glycan elongation. By comparison, the mimivirus glycosyltransferase candidate L137 predominantly behaved as a monomeric species under the tested conditions.

These findings demonstrate that multidomain glycosyltransferases employ diverse dynamic organizational strategies ranging from rigid recognition architectures to highly flexible and reversible catalytic assemblies. Our results further suggest that glycosyltransferase function is governed not only by catalytic-domain structure, but also by dynamic conformational coordination adapted to distinct catalytic demands.

## Introduction

Many glycosyltransferases possess multidomain architectures that extend beyond catalytic function alone [1]. These enzymes frequently contain auxiliary domains, flexible linker regions, oligomerization interfaces, or multiple catalytic modules that contribute to substrate recognition, catalytic coordination, and higher-order molecular organization during glycan biosynthesis. Glycosyltransferases generate the structural diversity of glycans by transferring monosaccharides from activated donor substrates to specific acceptor molecules and thereby play essential roles in protein quality control, extracellular matrix organization, cell–cell communication, and polysaccharide biosynthesis [2]. Recent work has provided important insights into processivity, substrate retention, and chain-length control in several glycosyltransferases [3]. However, relatively little is known about how multidomain glycosyltransferases are structurally organized and dynamically behave in solution. The dynamic organization of multidomain glycosyltransferases may be particularly important for enzymes involved in glycan elongation. Unlike glycosyltransferases that catalyze a single transfer reaction, elongation-related enzymes must repeatedly engage a growing glycan chain, position the non-reducing terminus, and coordinate sequential transfer reactions during polymer synthesis. Some elongating glycosyltransferases contain two distinct catalytic activities within a single polypeptide chain and must repeatedly reposition catalytic domains while accommodating elongating glycans during repeated catalytic cycles. Thus, understanding how catalytic domains are dynamically organized in solution is likely essential for clarifying the molecular basis of glycan elongation and polysaccharide synthesis.

Recent structural studies have provided important insights into the architectures of multidomain glycosyltransferases. POMGNT2, an O-mannosyl glycan biosynthetic enzyme required for core M3 formation on α-dystroglycan, forms a rigid dimeric structure in which an O-mannosylated acceptor peptide is recognized between two protomers through cooperative interactions involving catalytic and fibronectin type III-like domains [4–6]. In contrast, elongation-related glycosyltransferases such as LARGE1 and K4CP are thought to require more dynamic coordination of catalytic domains during polymer synthesis. LARGE1, a bifunctional glycosyltransferase responsible for matriglycan elongation, contains xylosyltransferase and glucuronyltransferase domains that support alternating transfer reactions during polymerization [7,8]. Structural analyses suggested that LARGE1 dimerization contributes to catalytic-domain organization, whereas recent enzymatic studies demonstrated processive matriglycan elongation on dystroglycan substrates [9,10]. K4CP, a bacterial chondroitin polymerase, also contains two catalytic domains responsible for alternating transfer of N-acetylgalactosamine and glucuronic acid during chondroitin elongation [11,12]. In addition, giant viruses such as Mimivirus encode glycosyltransferase-like proteins associated with biosynthesis of long glycans decorating viral fibers, suggesting that dynamic glycan assembly systems are not restricted to mammalian or bacterial pathways [13].

Despite these advances, static crystal structures and cryo-electron microscopy reconstructions do not necessarily reveal how glycosyltransferases behave in solution or how catalytic domains dynamically reorganize during substrate engagement and glycan elongation. Weak oligomerization, reversible assembly, large-amplitude interdomain motion, and substrate-dependent conformational changes may be difficult to capture using static structural methods alone. High-speed atomic force microscopy (HS-AFM) is particularly well suited for addressing this problem because it enables direct visualization of individual protein particles and their conformational dynamics in solution. When integrated with complementary solution biophysical approaches such as analytical ultracentrifugation (AUC), mass photometry, small-angle X-ray scattering (SAXS), and native mass spectrometry, HS-AFM provides a powerful framework for linking static structural models with dynamic molecular behavior.

In the present study, we comparatively investigated the assembly states and conformational dynamics of four multidomain glycosyltransferases—POMGNT2, LARGE1, K4CP, and L137—using HS-AFM integrated with complementary structural and solution biophysical approaches. By applying HS-AFM as a common single-molecule imaging platform, we aimed to clarify how structurally distinct glycosyltransferases employ different modes of catalytic-domain organization and conformational coordination in solution.

## Results and Discussion

### POMGNT2 forms a rigid dimeric reference architecture

POMGNT2 is a glycosyltransferase involved in the biosynthesis of the core M3 O-mannosyl glycan on α-dystroglycan and catalyzes the transfer of N-acetylglucosamine to O-mannosylated substrates. Previous crystallographic studies demonstrated that POMGNT2 forms a stable dimeric structure in which the acceptor peptide is recognized through cooperative interactions involving both catalytic and fibronectin type III-like (FnIII) domains from two protomers, suggesting that dimerization contributes directly to substrate recognition and catalytic specificity rather than merely stabilizing the protein structure.

We independently determined the crystal structure of the luminal region of POMGNT2 in the presence of UDP (PDB code: 8KB7) (Fig. 2A). The structure revealed a multidomain architecture consisting of an N-terminal catalytic domain and a C-terminal FnIII domain and closely resembled the previously reported dimeric organization of POMGNT2 (Supplementary Fig. S1). In this architecture, an O-mannosylated acceptor peptide is recognized by a composite binding site formed between two protomers, supporting a structurally constrained mode of site-selective substrate recognition.

**Fig. 1.**
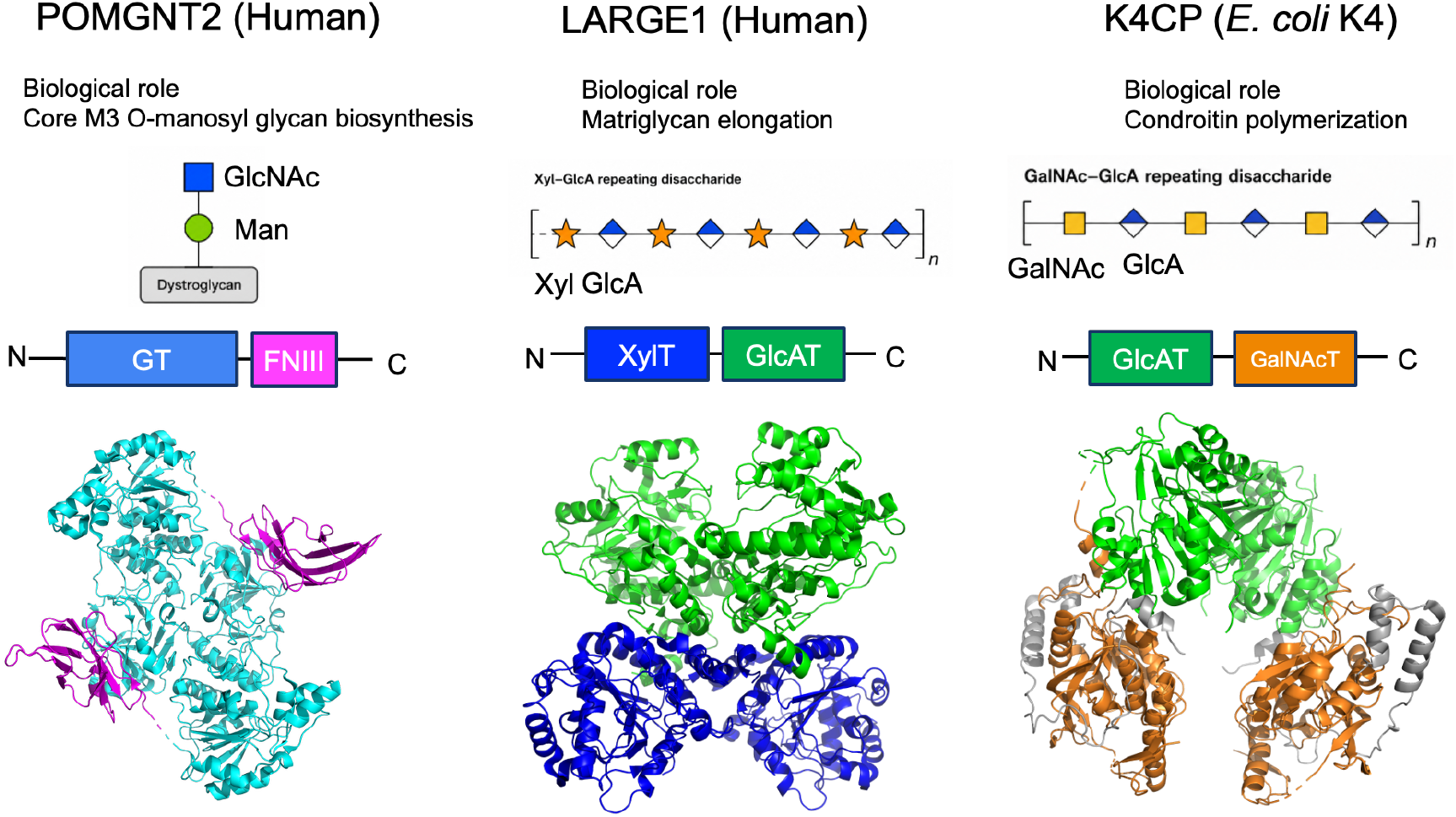
Domain architectures and dimeric structures of POMGNT2, LARGE, and K4CP. Domain organization, representative glycan products, and dimeric structures of POMGNT2, LARGE, and K4CP are shown. POMGNT2 consists of an N-terminal glycosyltransferase domain and a C-terminal FnIII domain and transfers GlcNAc to O-mannosylated glycans. Its dimeric crystal structure is shown below the domain schematic (PDB code: 7E9J). LARGE1 consists of an N-terminal xylosyltransferase domain (blue) and a C-terminal glucuronyltransferase domain (green), which together catalyze the alternating addition of xylose and glucuronic acid to generate matriglycan. Its dimeric structure is shown based on the cryo-EM structure (PDB code: 7ui7). K4CP is a bacterial chondroitin polymerase consisting of an N-terminal glucuronyltransferase domain (green) and a C-terminal N-acetylgalactosaminyltransferase domain (orenge), which catalyze the alternating transfer of glucuronic acid and N-acetylgalactosamine during chondroitin chain elongation. Its dimeric X-ray structure is shown below the domain schematic (PDB code: 2Z87).

**Fig. 2.**
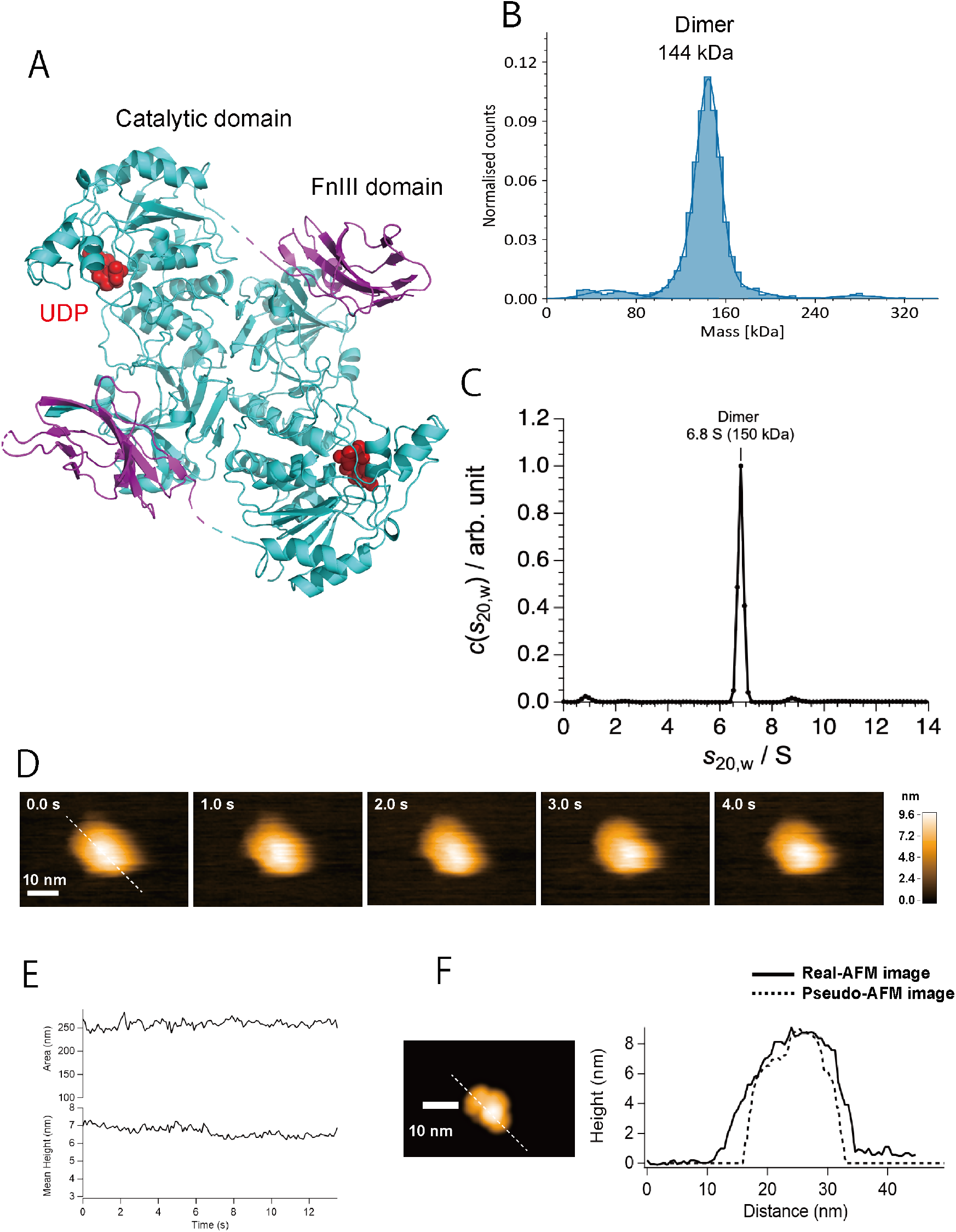
Structural and solution-state characterization of POMGNT2. (A) Crystal structure of the luminal region of POMGNT2 in complex with UDP. The structure shows a dimeric architecture composed of an N-terminal catalytic domain and a C-terminal fibronectin type III-like domain. The two protomers are shown in green and cyan. (B) Mass photometry analysis of recombinant POMGNT2. The dominant mass peak corresponds to the dimeric species, indicating that POMGNT2 predominantly exists as a dimer in solution under the AFM conditions. (C) Analytical ultracentrifugation analysis of recombinant POMGNT2. The sedimentation profile supports stable dimer formation in solution. (D) Representative high-speed AFM images of POMGNT2. Time-lapse images show relatively uniform particles with limited large-scale conformational fluctuation. Scale bars, 10 nm. Imaging conditions: 106 × 67 pixels, 56 × 38 nm^2^ scan area, 100 ms per frame; height scale, 0–9.6 nm. (E) Time-dependent changes in particle height and projected area measured from HS-AFM images. POMGNT2 particles exhibited only minor fluctuations during observation, indicating a comparatively rigid architecture. (F) Simulated AFM image and cross-sectional height profile of the POMGNT2 dimer generated from the crystal structure. The simulated AFM model recapitulated the overall morphology and height profile of the experimentally observed HS-AFM particles, supporting the conclusion that POMGNT2 maintains a crystallographic dimer-like architecture in solution.

To examine whether this dimeric organization is maintained in solution, we analyzed recombinant POMGNT2 using mass photometry, AUC, and HS-AFM (Fig. 2B, 2C, and 2D, respectively). Mass photometry showed a dominant particle population corresponding to the dimeric species, while analytical ultracentrifugation similarly supported stable dimer formation under the tested conditions.

HS-AFM imaging revealed relatively uniform particles with limited large-scale conformational fluctuation compared with the elongation-related glycosyltransferases analyzed in this study (Fig. 2E). The observed particle dimensions were generally consistent with the crystallographic model, indicating that the rigid dimeric organization of POMGNT2 is maintained both in crystal structures and in solution (Fig. 2F).

The relatively constrained organization of POMGNT2 contrasts with the more dynamic and heterogeneous behaviors observed for elongation-related glycosyltransferases such as LARGE1 and K4CP. Unlike polymerizing glycosyltransferases that repeatedly reorganize catalytic domains during sequential transfer reactions, POMGNT2 catalyzes a site-selective transfer reaction on a defined glycopeptide substrate. In this system, dimerization appears to be structurally required for formation of a composite substrate-recognition platform spanning two protomers. The relatively rigid organization of POMGNT2 may therefore stabilize precise positioning of the O-mannosylated glycopeptide acceptor during catalysis rather than support dynamic catalytic-domain rearrangement. These observations establish POMGNT2 as a useful reference architecture for comparing the diverse dynamic organizational strategies of multidomain glycosyltransferases.

### LARGE1 exhibits heterogeneous and reversible assembly behavior

LARGE1 is a bifunctional glycosyltransferase responsible for matriglycan elongation on α-dystroglycan through alternating xylosyltransferase and glucuronyltransferase reactions. Previous structural studies demonstrated that LARGE1 forms a dimeric architecture in which catalytic domains from opposing protomers are spatially arranged to support sequential glycan elongation. In addition, recent enzymatic reconstitution studies showed that LARGE1 processively polymerizes length-controlled matriglycan on dystroglycan substrates, raising the question of how catalytic-domain organization is dynamically coordinated during repeated transfer reactions (Fig. 3A).

**Fig. 3.**
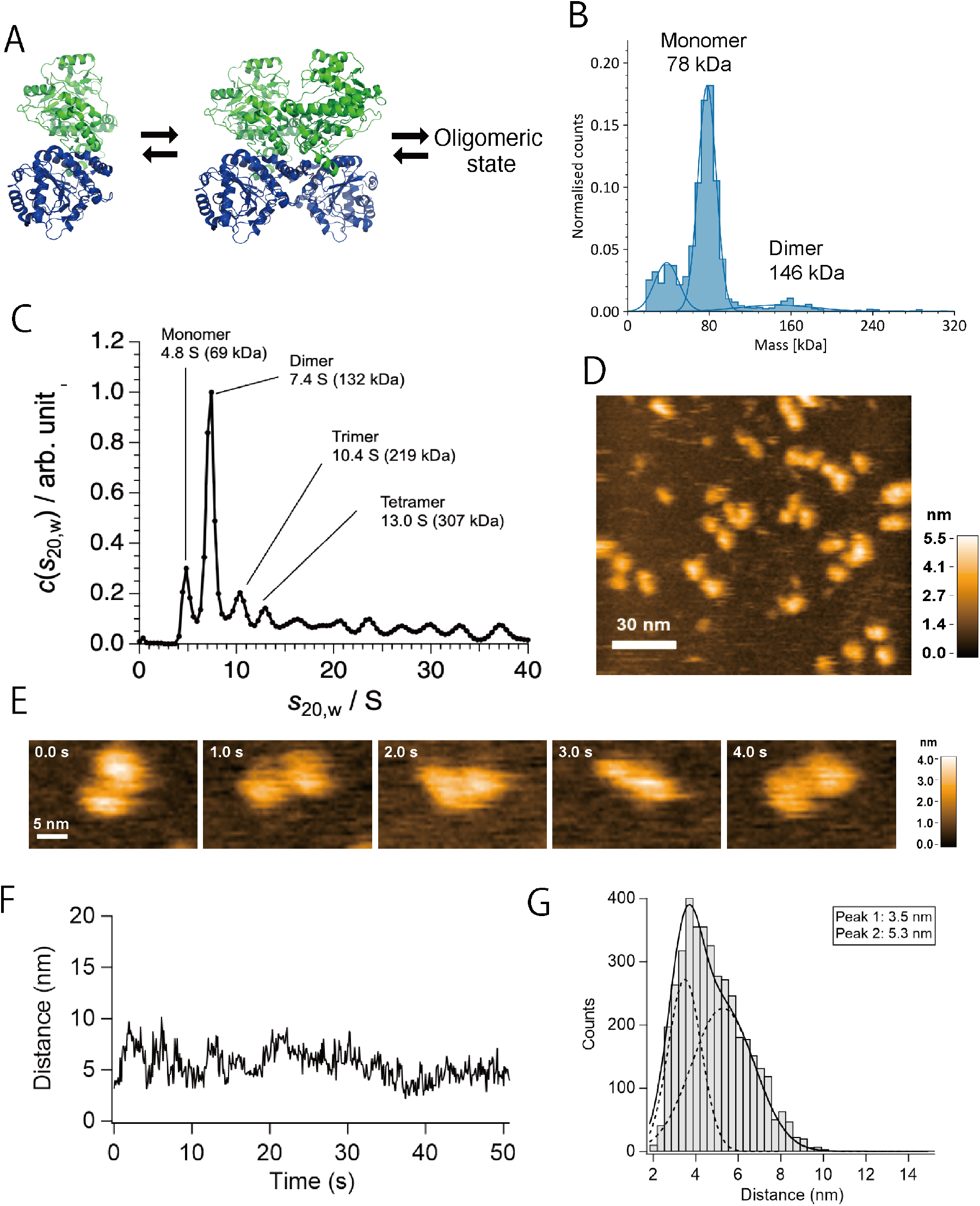
LARGE1 exhibits heterogeneous and reversible assembly behavior in solution. (A) Schematic representation of LARGE1-mediated matriglycan elongation on α-dystroglycan. LARGE1 catalyzes alternating xylosyltransferase and glucuronyltransferase reactions, and previous structural studies proposed a dimeric arrangement that may support coordinated, processive glycan elongation. (B) Mass photometry analysis of recombinant LARGE1. The mass distribution showed heterogeneous particle populations, including species compatible with monomer-like and dimer-like assemblies. Smaller particles became more prominent under dilute conditions, suggesting concentration-dependent self-association. (C) AUC profile of LARGE1. The broad and polydisperse weight concentration distribution contained species consistent with monomeric, dimeric, and higher-order assemblies, indicating that LARGE1 does not adopt a single dominant oligomeric state under the conditions tested. (D) Representative HS-AFM image of LARGE1 particles. Particles showed heterogeneous apparent sizes and shapes, with smaller monomer-like particles and larger dimer-like or oligomer-like assemblies observed in the same field. Scale bar, 30 nm; height scale, 0–5.5 nm. (E) Time-lapse HS-AFM images of a representative single LARGE1 particle. The two catalytic domains were resolved within an individual monomeric LARGE1 particle, and their relative positions fluctuated over time, indicating flexible interdomain motion. Imaging conditions: 54 × 32 pixels, 28 × 18 nm^2^ scan area, 100 ms per frame; scale bar, 5 nm; height scale, 0–4.0 nm. (F) Time-dependent distance change between the two domains within a representative single LARGE1 particle. (G) Distance distribution between the two domains measured from time-lapse HS-AFM images. The broad distribution supports dynamic fluctuation of the interdomain arrangement within individual LARGE1 particles. The distribution comprised two populations centered at ∼3.5 and 5.3 nm (n = 4,066 measurements from 6 particles).

To examine the assembly behavior of LARGE1 under solution conditions, we analyzed a soluble LARGE1 construct lacking the cytoplasmic, transmembrane, and stem (CTS) regions, using mass photometry, analytical ultracentrifugation, and HS-AFM. Mass photometry detected heterogeneous particle populations corresponding to both monomer-like and dimer-like species, with smaller particles becoming more prominent under dilute conditions (Fig. 3B). AUC similarly showed a broad and polydisperse sedimentation profile containing species compatible with monomeric, dimeric, and higher-order assemblies (Fig. 3C). Because of this heterogeneity, the apparent molecular masses were interpreted cautiously and did not indicate a single dominant oligomeric state under the present experimental conditions.

HS-AFM imaging further revealed heterogeneous particle morphologies and dynamic behavior. At the single-molecule level, individual LARGE1 particles displayed two discernible domains connected in a flexible arrangement, with the relative positions of these domains fluctuating over time (Fig. 3E, 3F, 3G). Some particles exhibited dimensions compatible with smaller monomer-like species, whereas others appeared larger and were consistent with dimer-like or oligomeric assemblies (Fig. 3D). Several particles also exhibited transient domain rearrangements and fluctuating interdomain organization. The interdomain distance distribution comprised two populations centered at approximately 3.5 nm (compact) and 5.3 nm (extended) (n = 4,066 measurements from 6 particles). The two lobes resolved within individual particles correspond to the two catalytic domains (xylosyltransferase and glucuronyltransferase) of a single LARGE1 molecule, and the interdomain distance analysis (Fig. 3F, 3G) was performed on such monomeric particles. Compared with the relatively rigid and uniform organization observed for POMGNT2, LARGE1 displayed substantially greater heterogeneity in both apparent particle size and dynamic behavior.

These observations suggest that the soluble LARGE1 construct does not behave as a rigidly fixed dimer in solution under dilute conditions. Rather than representing a single stable assembly state, LARGE1 appears to undergo reversible and condition-dependent self-association in solution. These observations likely reflect the dynamic behavior of the soluble catalytic construct outside the native Golgi membrane environment, where membrane confinement and local molecular crowding may contribute to stabilization of productive catalytic assemblies. Interactions with substrate glycoproteins may also help organize catalytic-domain positioning during matriglycan elongation.

The heterogeneous and reversible assembly behavior of LARGE1 contrasts with the rigid dimeric organization of POMGNT2 and suggests that elongation-related glycosyltransferases may employ more flexible organizational strategies to support repeated glycosyltransfer reactions. At the same time, the observed behavior of LARGE1 was less dynamically pronounced than that of K4CP, which exhibited large-amplitude interdomain motion and substrate-responsive conformational reorganization, as described below.

### K4CP undergoes dynamic catalytic reorganization through weak dimerization and large-amplitude domain motion

K4CP is a bacterial chondroitin polymerase that catalyzes the alternating transfer of N-acetylgalactosamine and glucuronic acid during chondroitin chain elongation. Structural studies previously showed that K4CP contains two catalytic domains responsible for the two glycosyltransferase activities required for polymer synthesis. Unlike POMGNT2, which catalyzes a site-selective transfer reaction within a relatively rigid dimeric architecture, K4CP repeatedly coordinates alternating catalytic reactions while accommodating an elongating polysaccharide chain, suggesting a greater requirement for dynamic catalytic-domain organization.

To investigate the assembly state of K4CP in solution, we performed mass photometry, AUC, and native mass spectrometry analyses. Mass photometry detected both monomeric and dimeric particle populations, with the monomeric species predominating under dilute conditions and the dimeric population increasing at higher protein concentrations (Fig. 4B). AUC similarly revealed concentration-dependent sedimentation behavior consistent with weak and reversible dimerization (Fig. 4C). Deconvoluted masses were close to the theoretical masses expected for monomeric and dimeric K4CP, supporting the assignment of the observed ion series. Although native MS ion intensities should not be interpreted as direct solution abundances, the detection of both species was consistent with the concentration-dependent reversible self-association indicated by mass photometry and AUC. (Fig. 4D). Together, these observations indicate that K4CP does not form a constitutively stable oligomeric assembly but instead undergoes concentration-dependent and reversible self-association. To further examine the solution organization of K4CP, we performed SAXS analysis (Fig. 4E and Supplementary Fig. S2). The experimental scattering profile at 4.0 mg/mL was consistent with dimer-compatible structural models than with a compact monomeric arrangement alone (*χ*^2^ = 105 and 3.4 for monomer and dimer, respectively).

**Fig. 4.**
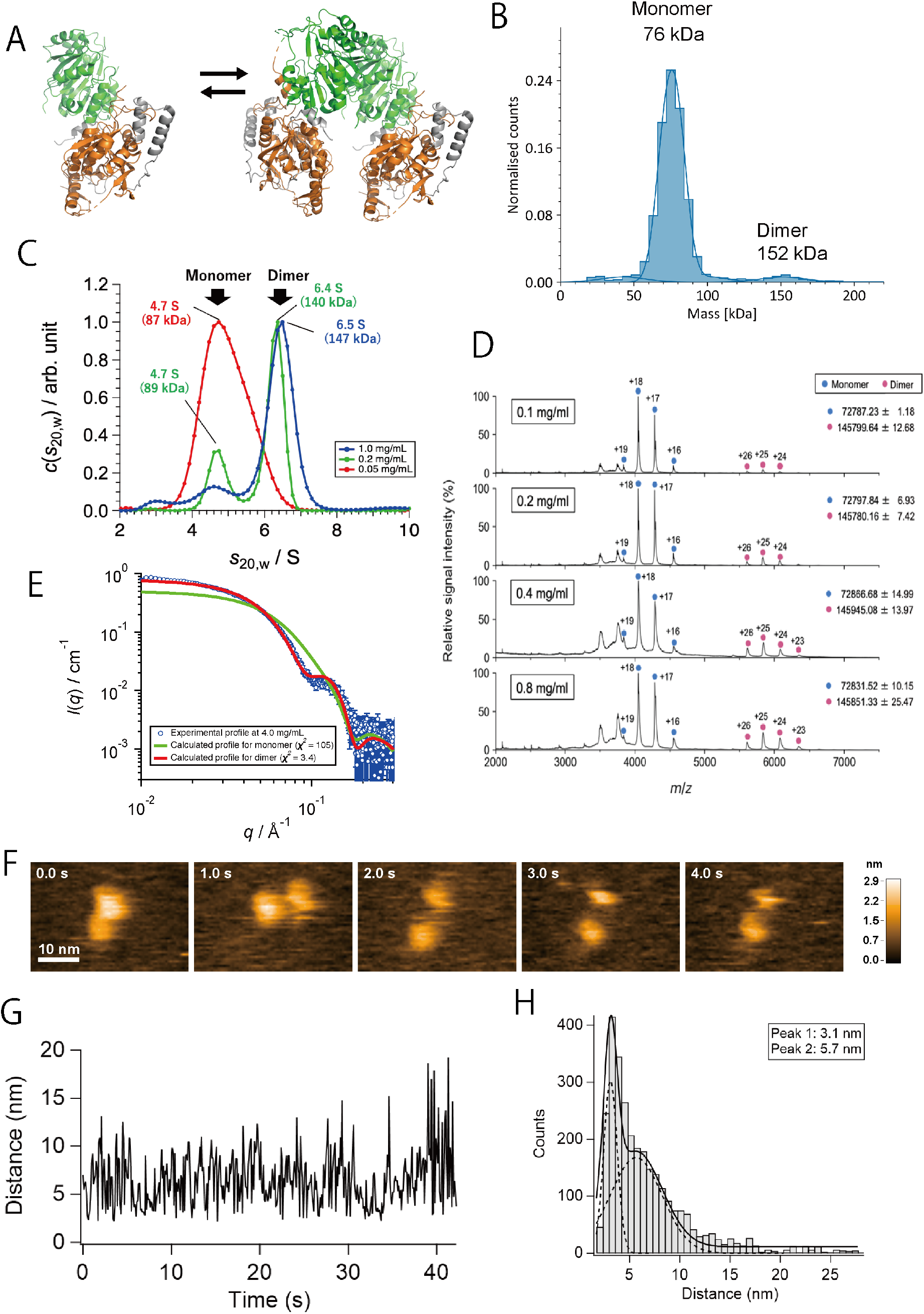
K4CP displays reversible self-association and pronounced interdomain dynamics during chondroitin polymerization. (A) Structural models of K4CP. The monomer contains two catalytic domains responsible for the alternating transfer of N-acetylgalactosamine and glucuronic acid during chondroitin chain elongation. The dimer model was generated from the crystal structure and illustrates the crystallographic dimeric arrangement of K4CP. (B) Mass photometry analysis of recombinant K4CP. The mass distribution revealed a predominant monomeric population together with a smaller dimeric population, indicating heterogeneous assembly in solution. (C) AUC profiles of K4CP at different protein concentrations. Red, green, and blue circles mean the prifle for 0.05, 0.2, and 1.0 mg/mL, respectively. The weight concentration distributions showed a shift from monomer-dominant species at lower concentration toward dimer-compatible species at higher concentration, supporting weak and reversible dimerization.(D)Native mass spectrometry analysis of K4CP. The dimer-associated signals became more evident at higher protein concentrations, supporting the presence of concentration-dependent self-association. (E)SAXS analysis of K4CP. Blue circles, green line, and red line represent the experimental profile, calculated profile for monomer, calculated profile for dimer, respectively, generated from the crystal structure of K4CP (PDB ID: 2Z87). The experimental scattering profile at 4.0 mg/mL was compared with the dimer model generated from the crystal structure, and the fitting supported the presence of dimer-compatible K4CP species in solution. (F) Time-lapse HS-AFM images of a representative K4CP particle. Individual particles exhibited repeated changes in the relative positions of two lobular domains, revealing large-amplitude interdomain motion at the single-molecule level. Imaging conditions: 75 × 48 pixels, 39 × 27 nm^2^ scan area, 100 ms per frame; scale bar, 10 nm; height scale, 0–2.9 nm. (G) Time-dependent distance change between two domains within a representative single K4CP particle. (H) Distribution of interdomain distances measured from HS-AFM movies. The broad distribution indicates that K4CP samples multiple conformational states, including compact and extended arrangements. The two populations were centered at ∼3.1 and 5.7 nm (n = 2,971 measurements from 5 particles).

HS-AFM imaging directly revealed pronounced conformational dynamics of K4CP in solution (Fig. 4F-4H). Individual particles frequently underwent repeated open–closed transitions accompanied by large-amplitude interdomain motion. Distance measurements between lobular domains demonstrated broad conformational distributions, with compact and extended populations centered at approximately 3.1 and 5.7 nm, respectively, and transient excursions to interdomain separations of up to ∼20 nm (Fig. 4G, 4H) Transient transitions between compact and extended conformations were repeatedly observed within individual particles, indicating highly dynamic catalytic-domain rearrangement in solution. Compared with POMGNT2 and LARGE1, K4CP displayed substantially greater conformational flexibility and dynamic rearrangement behavior.

To investigate whether substrate binding influences K4CP organization, we analyzed the protein in the presence of chondroitin oligosaccharides. Under these conditions, HS-AFM imaging suggested a tendency toward more compact conformations with reduced interdomain separation, although this effect was not quantified in the present study. These preliminary observations are consistent with the possibility that catalytic-domain organization in K4CP may be modulated by substrate binding during polysaccharide elongation. Such large-amplitude and substrate-responsive motions may facilitate repeated repositioning of catalytic sites during sequential glycan transfer reactions.

The highly dynamic behavior of K4CP contrasts strongly with the rigid dimeric organization of POMGNT2 and extends the heterogeneous and reversible assembly behavior observed for LARGE1. Whereas static crystal structures previously provided important architectural snapshots of multidomain glycosyltransferases, the present HS-AFM analyses demonstrate that K4CP exists as a highly dynamic molecular assembly in solution. These observations suggest that flexible catalytic-domain organization represents an important strategy for coordinating alternating glycosyltransfer reactions during polysaccharide elongation.

### Comparative dynamic organization of multidomain glycosyltransferases

Structural and dynamic comparisons of POMGNT2, LARGE1, and K4CP revealed distinct modes of molecular assembly among multidomain glycosyltransferases. Although all three enzymes contain multiple structural modules involved in substrate recognition and catalysis, their assembly behaviors and conformational dynamics differed substantially in solution. POMGNT2 maintained a relatively rigid dimeric architecture with limited large-scale conformational fluctuation, whereas LARGE1 exhibited heterogeneous and reversible assembly behavior under dilute solution conditions. In contrast, K4CP displayed pronounced conformational flexibility characterized by large-amplitude interdomain motion and substrate-responsive structural rearrangement.

The observed diversity in solution behavior appears to correlate with the distinct catalytic demands imposed on each enzyme system. POMGNT2 catalyzes a site-selective transfer reaction on a defined glycopeptide substrate during core M3 glycan biosynthesis, whereas LARGE1 and K4CP repeatedly coordinate alternating glycosyltransfer reactions during glycan elongation and polysaccharide synthesis. In POMGNT2, dimerization appears structurally linked to formation of a composite substrate-recognition platform, whereas LARGE1 and K4CP appear to prioritize more flexible catalytic-domain coordination during repeated transfer reactions. Previous structural studies suggested relatively compact catalytic-domain arrangements for both LARGE1 and K4CP. However, the present HS-AFM analyses demonstrated that these multidomain glycosyltransferases exhibit substantially broader conformational distributions in solution than implied by static structural models alone.

Comparison of catalytic-domain distances further highlighted these differences in conformational coordination. The experimentally observed distance distributions obtained by HS-AFM extended beyond those predicted from previously reported crystallographic models, particularly in the case of K4CP. These observations indicate that multidomain glycosyltransferases exist as dynamic conformational ensembles in solution rather than as rigid molecular assemblies fixed in a single structural state.

In addition to the three structurally and dynamically characterized glycosyltransferases described above, preliminary analyses of the viral glycosyltransferase candidate L137 suggested that predominantly monomeric and flexible assemblies may also occur in glycosyltransferase systems outside canonical mammalian and bacterial pathways. Although the current characterization of L137 remains limited, these observations further expand the apparent spectrum of solution behaviors observed among multidomain glycosyltransferases (Supplementary Fig. S3).

Taken together, these comparative analyses suggest that multidomain glycosyltransferases can employ distinct organizational strategies ranging from rigid architectures to reversible assemblies and dynamic conformational cycling, depending on the structural and catalytic requirements of individual glycan synthesis systems.

**Table 1.**
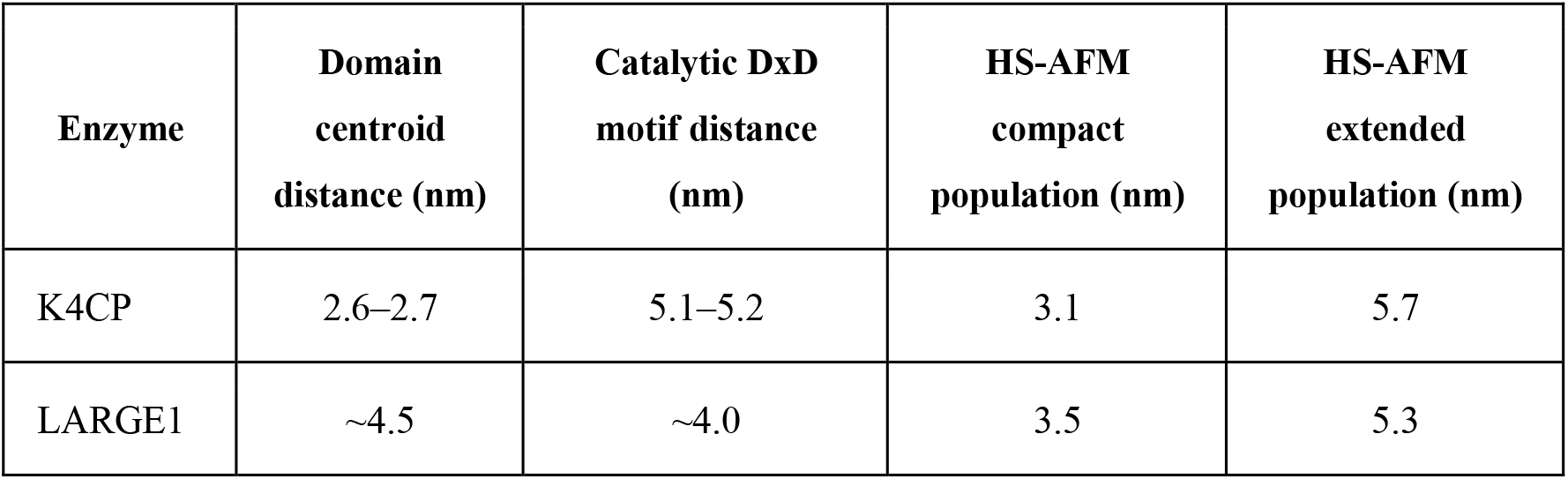
Comparison of model-derived and HS-AFM-derived interdomain distances in K4CP and LARGE1.

### Functional implications of dynamic catalytic-domain coordination

The present study revealed a broad spectrum of molecular assemblies and conformational behaviors among multidomain glycosyltransferases. Comparative analyses using crystallography, solution biophysics, and HS-AFM demonstrated that these enzymes range from relatively rigid dimeric assemblies to highly flexible and reversible catalytic systems. Importantly, the observed diversity was associated not simply with oligomeric state, but with distinct modes of catalytic-domain coordination and conformational regulation adapted to different catalytic demands.

Our results further highlight the importance of integrating static structural biology with direct analyses of solution dynamics. Crystal structures and cryo-electron microscopy reconstructions provide essential architectural snapshots of glycosyltransferases and can reveal distinct conformational states. However, because these approaches do not directly follow time-dependent molecular motions in solution, complementary methods are needed to characterize transient conformational fluctuations, reversible assembly behavior, and substrate-responsive rearrangements. By combining HS-AFM with complementary solution biophysical methods, including analytical ultracentrifugation, mass photometry, small-angle X-ray scattering, and native mass spectrometry, we directly visualized dynamic molecular behaviors extending beyond static structural models.

The contrasting behaviors observed for POMGNT2, LARGE1, and K4CP suggest that multidomain glycosyltransferases employ distinct structural solutions to accommodate different catalytic requirements. In POMGNT2, the rigid dimeric architecture appears structurally linked to formation of a composite substrate-recognition platform spanning two protomers. Such a constrained arrangement may stabilize precise positioning of the O-mannosylated glycopeptide acceptor during catalysis and thereby support highly site-selective substrate recognition. In contrast, the more flexible and reversible assemblies observed for LARGE1 and K4CP may facilitate accommodation and repeated repositioning of elongating glycan chains during polymerization reactions.

In particular, the large-amplitude and substrate-responsive motions observed for K4CP indicate that multidomain glycosyltransferases can dynamically reorganize catalytic modules in response to functional demands. Rather than behaving as rigid molecular machines fixed in a single architecture, elongation-related glycosyltransferases may sample multiple conformational states during repeated catalytic cycles. The heterogeneous behavior observed for LARGE1 further suggests that productive catalytic assemblies may be stabilized by membrane confinement, local molecular crowding, or substrate interactions within the native Golgi environment.

The exploratory observations obtained for the viral glycosyltransferase candidate L137 further support the idea that glycosyltransferases can adopt structurally diverse solution states beyond stable oligomeric assemblies. Although additional structural and functional analyses will be required to clarify the biological significance of L137 organization, its predominantly monomeric behavior expands the apparent range of molecular assemblies accessible to glycosyltransferase systems.

Together, these observations establish HS-AFM as a powerful platform for investigating the dynamic structural biology of glycosyltransferases and other multidomain enzymes. Future studies combining HS-AFM with substrate-bound and membrane-associated glycosyltransferase systems may further clarify how catalytic domains are dynamically coordinated during glycan elongation and polysaccharide synthesis. More broadly, our comparative analyses suggest that multidomain glycosyltransferases employ distinct organizational strategies—including structural constraint, reversible assembly, and conformational cycling—to meet different catalytic demands. Dynamic solution organization therefore represents an additional layer of functional regulation that complements catalytic-domain architecture and substrate specificity in glycan biosynthetic systems.

## Materials and Methods

### Expression and purification of recombinant glycosyltransferases

DNA fragments encoding soluble human LARGE1 residues 131–756 and human POMGNT2 residues 52–580 were cloned into the p3FLAG-CMV-9 vector (Sigma) for expression as N-terminal FLAG-tagged proteins. The resulting plasmids were transiently transfected into Expi293F cells according to the manufacturer’s instructions, and recombinant proteins were expressed in suspension culture. Culture supernatants were collected and applied to anti-FLAG M2 affinity resin (Sigma). Bound proteins were eluted with 20 mM glycine buffer (pH 2.5) and further purified by size-exclusion chromatography using a HiLoad 16/600 Superdex 200 column (Cytiva) equilibrated with 20 mM Tris-HCl and 150 mM NaCl. Fractions containing the target proteins were pooled and used for subsequent biochemical and structural analyses.

For L137, a DNA fragment encoding the target region was cloned into the pBAD vector for expression as an N-terminal His-tagged protein. The plasmid was purchased from Thermo Fisher Scientific and transformed into Escherichia coli DH5α. Recombinant protein expression was induced in bacterial culture with 0.2% L-arabinose. Cells were harvested by centrifugation and lysed, and the clarified lysate was applied to a Ni^2+^-affinity column for purification of the His-tagged protein. Fractions containing L137 were further purified by size-exclusion chromatography using a HiLoad 16/600 Superdex 200 column equilibrated with 20 mM Tris-HCl and 150 mM NaCl.

A DNA fragment encoding residues 58–686 of K4CP was synthesized by Fasmac and cloned into the pCold vector for expression as an N-terminal His-tagged protein. The plasmid was transformed into E. coli BL21-CodonPlus cells, and protein expression was induced with 0.3 mM IPTG at 18°C for approximately 20 h. Cells were harvested, lysed by sonication, and the clarified lysate was applied to a Ni^2+^-affinity column. Bound proteins were eluted with imidazole, and fractions containing His-tagged K4CP were concentrated and further purified by size-exclusion chromatography on a HiLoad 16/600 Superdex 200 column equilibrated with 20 mM Tris-HCl and 150 mM NaCl.

### Crystallization, data collection, and structure determination of POMGNT2

To obtain high-quality crystals, purified POMGNT2 was treated with sialidase and galactosidase before crystallization. The deglycosylated POMGNT2 protein was concentrated to 5 mg/mL in PBS containing 5 mM MnCl_2_, 5 mM UDP, and 2 mM mannosyl peptide (IHAT(Man)PTPV). Crystals were obtained after incubation at 20°C for 1 week in a reservoir solution containing 12% PEG 4000, 0.1 M trisodium citrate, pH 6.0, and 0.1 M sodium chloride.

For heavy-atom derivatization, native POMGNT2 crystals were soaked for 12 h in crystallization buffer supplemented with 10 mM K_2_PtCl_4_ using the Heavy Atom Screen Pt kit (Hampton Research). All crystals were cryoprotected with crystallization mother liquor supplemented with 15% PEG 400 and flash-cooled in liquid nitrogen.

Native and anomalous diffraction datasets were collected at BL44XU, SPring-8, at wavelengths of 0.9000 Å and 1.0717 Å, respectively. The native and Pt-derivatized POMGNT2 crystals belonged to space group P2_1_2_1_2_1_ and diffracted to resolutions of 2.80 Å and 3.71 Å, respectively. All diffraction data were processed using XDS [14]. The crystallographic parameters are summarized in Supplementary Table 1.

The structure of POMGNT2 was determined by the single-wavelength anomalous dispersion method using the 3.71 Å Pt-derivatized dataset. Initial phases were obtained with AutoSol in the PHENIX suite [15]. Because the Pt-derivatized and native crystals were sufficiently isomorphous, the SAD phase information was transferred to the native dataset. After density modification and phase extension to 2.80 Å, the resulting electron density map was of sufficient quality for model building.

Four POMGNT2 molecules were present in the asymmetric unit. Partial models for two molecules were built automatically using ARP/wARP [16]. Manual model building was performed in COOT [17]. The remaining two molecules in the asymmetric unit were placed by molecular replacement using MOLREP[18], with the determined POMGNT2 N-terminal domain structure as the search model. Refinement was performed using REFMAC5 [19], and the stereochemical quality of the final model was assessed using MolProbity [20]. Refinement statistics are summarized in Supplementary Table 1. Molecular graphics were prepared using PyMOL (http://www.pymol.org/).

### Mass photometry

Mass photometry measurements were performed using a TwoMP (Refeyn) to determine the molecular mass distributions of POMGNT2, LARGE1, K4CP, and L137 under dilute solution conditions according to the manufacture’s instruction. Protein samples were diluted immediately before measurement and loaded on measurement chambers at the final concentrations (POMGNT2: 0.096 nM, LARGE1: 9.0 nM, K4CP: 0.23 nM, L137: 7.3 nM) following focus adjustments with sample buffer. Movies were recorded using AcquireMP software (Refeyn) and single-particle landing events were analyzed using DiscoverMP software (Refeyn). Molecular masses were assigned based on contrast-to-mass calibration with BSA (66 kDa and 132 kDa) and bovine thyroglobulin (670 kDa).

### Analytical ultracentrifugation (AUC)

AUC measurements were performed to analyze the solution assembly states of POMGNT2, LARGE1, and K4CP in the corresponding experimental buffers. The measurements were performed using ProteomeLab XL-I (Beckman Coulter) at 25°C. The samples were loaded into cells equipped with 12 mm optical path aluminums center piece and set in an An-60Ti rotor (Beckman Coulter). Sedimentation velocity analysis was performed using Rayleigh interference optics at the rotor speed of 60,000 rpm.

The weight-concentration distributions as a function of sedimentation coefficient *c*(*s*_20,w_) was obtained using SEDFIT software version 15.01c [21]. The sedimentation coefficient was normalized to be the value at 20 °C in pure water, *s*_20,w_. Because the sedimentation coefficients of oligomeric proteins are influenced by molecular weightmolecular shape, and reversible self-association, oligomeric assignments from AUC were interpreted as approximate and were evaluated together with mass photometry, native MS, and SAXS data. [22]

### SAXS analysis of K4CP

K4CP protein samples were prepared at concentrations of 0.5, 1.0, 2.0, and 4.0 mg/mL and subjected to SAXS analysis. SAXS measurements were performed using a small-angle X-ray scattering instrument NanoSTAR (Bruker) installed at the Australian Nuclear Science and Technology Organisation (ANSTO) at 25°C. The sample-detector distance was set at 1,070 mm. The radius of gyration (*R*_g_) and forward scattering intensity (*I*(0)) were determined Guinier fitting to the low-*q* region (*q* < 1.3/*R*_g_) of the experimentally obtained scattering curves, according to the following equations:

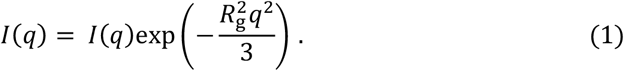

Here, *q* is a magnitude of scattering vector, *q* = (4π/*λ*)sin(*θ*/2), where *λ* and *θ* are wavelength of incident X-ray and scattering angle, respectively. Calculated scattering profiles based on the crystal structure were calculated using CRYSOL [23]. The agreement between the calculated SAXS profile *I*_cal_(*q*) and the experimental one *I*_exp_(*q*) was evaluated using the *χ*^2^-value in the following.

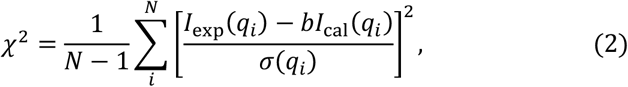

where *N* is the number of points in the scattering profile, *σ*(*q*) are the experimental errors, and *b* is the scaling factor as follows.

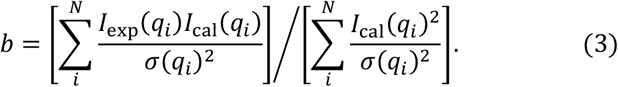

### Native mass spectrometry

K4CP protein samples were prepared at concentrations of 0.1, 0.2, 0.4, and 0.8 mg/mL and analyzed by native mass spectrometry (nMS). Prior to measurement, all samples were buffer-exchanged into 500 mM ammonium acetate, pH 8.0, using Bio-Spin 6 columns (Bio-Rad). The samples were loaded into gold-coated glass capillaries that had been adjusted to a final volume of approximately 2–5 μL and were immediately analyzed by nanoflow electrospray ionization mass spectrometry using a SYNAPT G2-*Si* HDMS mass spectrometer (Waters) in the positive ion mode.

Measurements were performed under the following instrument conditions: capillary voltage, 1.33 kV; sampling cone, 150 V; source offset, 150 V; nano flow gas, 0 bar; trap gas flow, 5 mL/min; trap collision energy, 0 V; transfer collision energy, 0 V; trap wave velocity, 12 m/s; transfer wave velocity, 12 m/s; and acquisition time, 5 min. The spectra were calibrated using cesium iodide dissolved in the solution of 50% 2-propanol and 50% water (*v/v*) at 1 mg/mL. Spectral data were analyzed using MassLynx software (Waters).

### High-speed atomic force microscopy

HS-AFM was used to visualize the morphology and conformational dynamics of POMGNT2, LARGE1 TM, K4CP, and L137. HS-AFM imaging was performed using a laboratory-built high-speed AFM [24] operated in tapping mode with a short cantilever (Olympus BL-AC10; nominal spring constant ∼0.1 N/m, resonance frequency ∼500 kHz in liquid, tip apex radius ∼2 nm). The free oscillation amplitude was 2–3 nm, and the set-point amplitude was 80–90% of the free amplitude. Stock proteins were diluted immediately before imaging and applied to freshly cleaved mica. After adsorption, the surface was gently rinsed with imaging buffer, and HS-AFM imaging was performed in liquid at room temperature. POMGNT2 (stock 0.285 mg/mL, 68 kDa, ∼4.2 µM) was diluted 1/100 to ∼42 nM in 20 mM Tris-HCl pH 7.5, 150 mM NaCl. The soluble LARGE1 construct (stock 0.012 mg/mL, 78 kDa, ∼0.15 µM) was imaged undiluted (∼0.15 µM) and at 1/5 dilution (∼30 nM) in 20 mM Tris-HCl pH 7.5, 150 mM NaCl. K4CP (stock 2 mg/mL, 75 kDa, ∼26.7 µM) was diluted 1/5000 to ∼5.3 nM in 50 mM Tris-HCl pH 8.0, 50 mM NaCl. L137 (stock 1.3 mg/mL, 78 kDa, ∼16.7 µM) was diluted 1/2100 to ∼8 nM in 20 mM Tris-HCl pH 7.5, 150 mM NaCl.

Time-lapse images were acquired with the pixel dimensions, scan areas, and frame rates given in the corresponding figure legends. For each enzyme, representative particles were selected for analysis, and apparent particle size, morphology, and domain-like movement were evaluated from sequential images. For K4CP, two domain-like lobes were identified, and the center-to-center distance between them was measured over time; interdomain distance distributions were compiled from 2,971 measurements across 5 particles (substrate-free condition); K4CP was additionally imaged in the presence of chondroitin oligosaccharides. For LARGE1, interdomain distance distributions were compiled from 4,066 measurements across 6 particles. For L137, inter-lobe angles among three lobular densities were measured, yielding populations centered near 73° and 112°.

Because HS-AFM observes surface-adsorbed particles, apparent molecular dimensions were interpreted with caution. Particle assignment as monomer-like or dimer-like was based on apparent size and morphology, but individual dynamic particles were not always assigned unambiguously to a specific oligomeric state. For POMGNT2, a simulated AFM image was generated from the crystal structure using in-house software (pyNuD Simulator) with a conical tip model (apex radius 2.0 nm, full cone angle 20°), low-pass filtered at a 2.0 nm cutoff to approximate the lateral resolution of HS-AFM, and compared with the experimental HS-AFM topography and cross-sectional height profile (Fig. 2F).

## Data analysis and statistics

Quantitative analyses of HS-AFM images were performed using in-house Python-based software (pyNuD). Interdomain distances were measured manually or semi-automatically from particle centers. Because tip–sample convolution broadens lateral features in AFM, absolute interdomain distances may be overestimated and were used primarily for relative comparison. Histograms were generated from pooled measurements obtained from multiple particles. Distance and angle histograms were fitted with Gaussian components to determine the population centers. For mass photometry and AUC, apparent molecular mass assignments were based on calibration or sedimentation analysis.

## Structural visualization

Structural models and molecular graphics were prepared using PyMOL. Previously reported structures of LARGE and K4CP were used for comparison with HS-AFM observations where appropriate. Structural models were aligned and displayed to illustrate domain organization, catalytic domain arrangement, and possible conformational changes.

## Conflicts of interest statement

The authors declared no conflicts of interest.

## Acknowledgements

We thank Ms. Kiyomi Senda and Ms. Kumiko Hattori (Nagoya City University) for their technical assistance. We are also grateful to Dr. Nobuo Sugiura (Aichi Medical University) and Dr. Koji Kimata (Aichi Medical University), and Dr. Yoshimitsu Kakuta (Kyushu University) for helpful discussions. We also thank Dr. Toshiya Kozai (Nagoya University), Mr. Naoya Masukane (Nagoya City University), and Ms. Saki Yoshida (Nagoya City University), whose affiliations are listed as those at the time of their contributions, for their contributions to the initial phase of this study. The synchrotron radiation experiments were performed at the BL41XU beamline of SPring-8 with the approval of the Japan Synchrotron Radiation Research Institute (JASRI) under Proposal Nos. 2021A6609 and 2022B6708. AUC measurements were performed under proposal numbers 30010 and 31006 at KURNS, Kyoto University. SAXS measurements were performed under proposal number 7005 at ANSTO.

## Funding

This work was supported in part by JST-CREST (grant number JPMJCR21E3 to K.K.), JST FOREST Program (grant number JPMJFR2255 to H.Y.), the Human Glycome Atlas Project, MEXT/JSPS KAKENHI (JP21H02625 and JP25K02410 to H.Y., JP19H03361 to T.S., and JP20K21495 and JP24H00599 to K.K.), the ExCELLS Advanced Co-creation Platform (Spatiotemporal atlas of dynamic structure and function of organelles, 23EXC601 to H.Y.), and ExCELLS “Golgi Atlas Project” (to K.K.) and Grant-in-Aid for Outstanding Research Group Support Program in Nagoya City University (Grant Number 2401101, 2520001 and 2530002), and by the Joint Research of the Exploratory Research Center on Life and Living Systems (ExCELLS) (ExCELLS programs No. 18-101 to T.U. and 22EXC601 to K.K. and T.U.). This work was also supported by KAKENHI (Grant Nos. JP24K01309 and JP26H01626 to T.U.) and the MEXT Promotion of Development of a Joint Usage/Research System Project: Coalition of Universities for Research Excellence Program (CURE) (Grant Number JPMXP1323015482). This work also utilized research equipment shared through the MEXT Project for promoting public utilization of advanced research infrastructure (Program for supporting construction of core facilities, JPMXS0441500026). This work also partialy supported by Platform Project for Supporting Drug Discovery and Life Science Research [Basis for Supporting Innovative Drug Discovery and Life Science Research (BINDS)] from AMED (award No. JP22ama121001j0001 to MS).

## Notes

### Competing Interest Statement

The authors have declared no competing interest.

