## Supplementary Figs. S1, S2 and S3, and Supplementary Table1 for "Dynamic organizational strategies of multidomain glycosyltransferases revealed by high-speed AFM and solution biophysics"

**Supplementary Fig. S1**

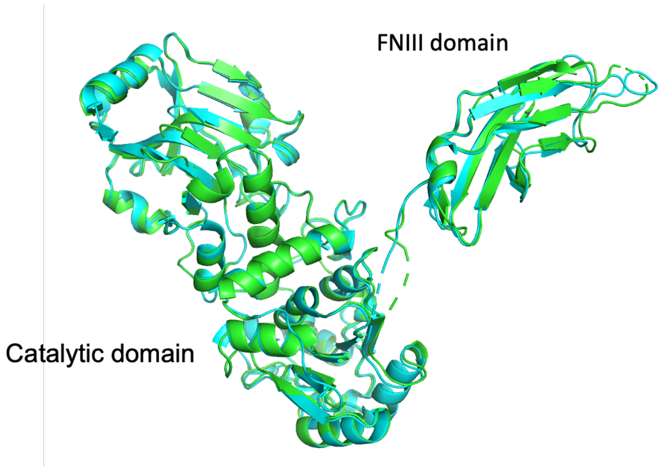


**Supplementary Fig. S1. Structural comparison of POMGNT2 with the previously reported structure.**

Superposition of the crystal structure of POMGNT2 determined in this study (green) (PDB code: 8KB7) with the previously reported POMGNT2 structure (cyan) (PDB code: 7E9J). POMGNT2 consists of an N-terminal catalytic domain and a C-terminal FnIII domain, which adopt a similar overall arrangement to that observed in the previous structure. Crystallographic data collection and refinement statistics are summarized in Supplementary Table 1.

**Supplementary Fig. S2**

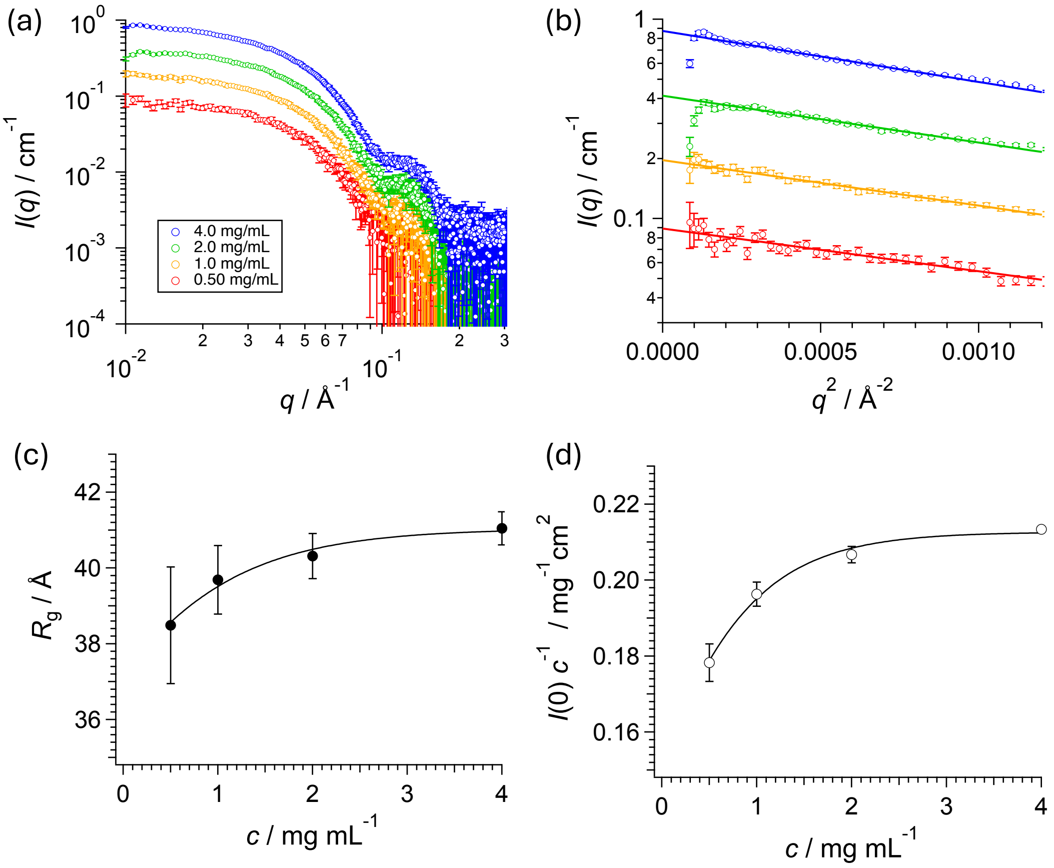


**Supplementary Fig. S2. SAXS analysis of K4CP at different protein concentrations.**

(a) SAXS profiles of K4CP were recorded at 0.5 (red circles), 1.0 (yellow circles), 2.0 (green circles), and 4.0 mg/mL (blue circles). (b) Guinier plots of the corresponding scattering intensities. Solid lines represent the mean square fitting by Guinier function. (c) Concentration dependence of the radius of gyrations $R_g$ offered by the Guinier analysis. Solid curve represent the fitting curve by Sigmoidal function for eye guide. (d) Concentration dependence of the forward scattering intensity $I(0)$ offered by the Guinier analysis. Solid curve represent the fitting curve by Sigmoidal function for eye guide. Both $R_g$ and $I(0)$ increased with the K4CP concentration and reached an apparent plateau at 4.0 mg/mL. This concentration dependentence indicates an increasing fraction of dimeric species, suggesting that the scattering contribution at 4.0 mg/mL is dominated by the dimer.

**Supplementary Fig. S3.**

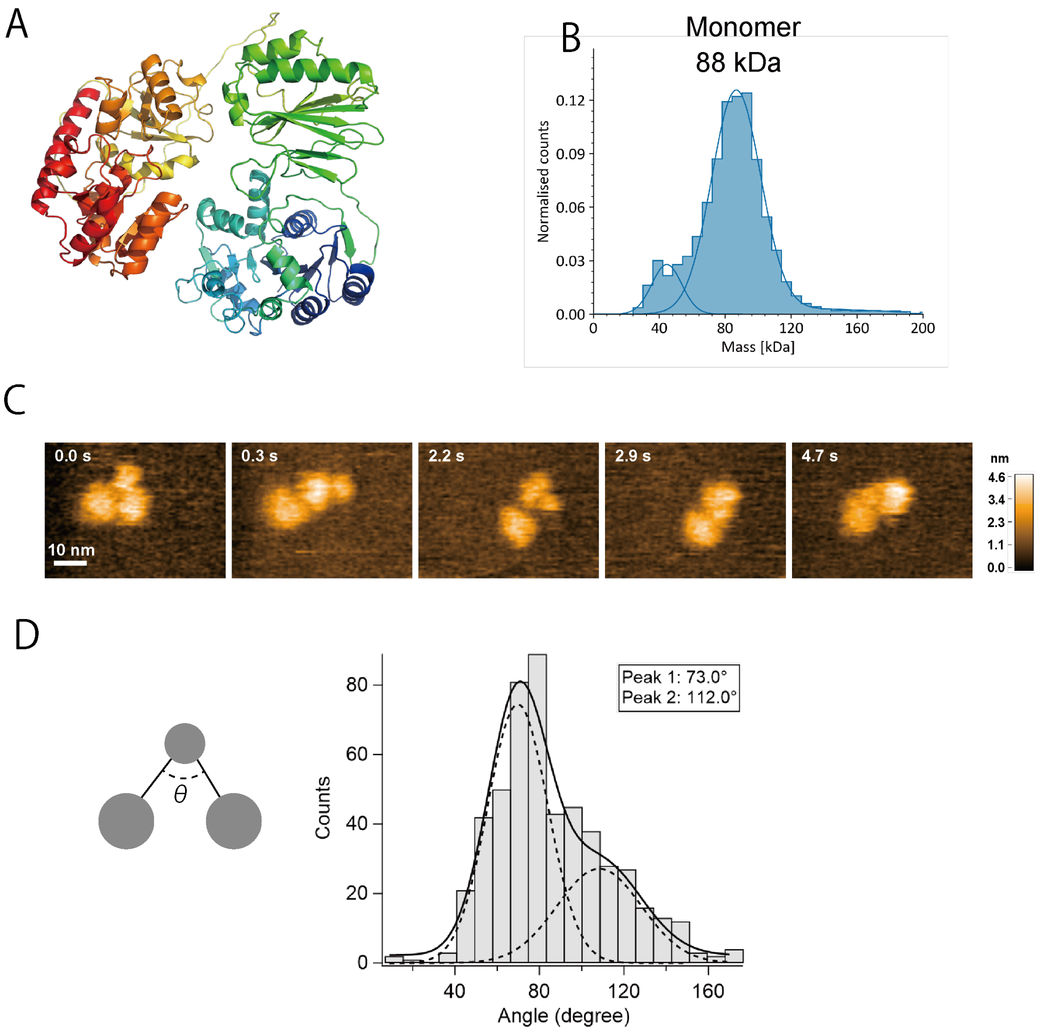


Supplementary Fig. S3. Structural and dynamic analysis of the giant-virus-derived L137 protein by mass photometry and HS-AFM.

(A) AF3-predicted structural model of giant-virus-derived L137 shown in a monomeric arrangement. (B) Mass photometry analysis of purified L137. The major population corresponded to monomeric L137. (C) Representative time-lapse HS-AFM images of L137 particles. Three lobular densities were repeatedly observed within individual particles, and their relative positions changed over time, indicating conformational flexibility. The two larger lobes likely correspond to the two domains of the L137 monomer, whereas the smaller central density represents an additional globular density between these domains, consistent with the AF3-predicted monomeric model. Imaging conditions: 93 × 64 pixels, 56 × 42 nm² scan area, 100 ms per frame; height scale, 0–4.6 nm. (D) Schematic representation of the three-lobed particle model used to quantify inter-lobe angles from HS-AFM images, together with the distribution of measured inter-lobe angles. The histogram was fitted with

Gaussian components, revealing a major population centered at approximately 73° and a minor population centered at approximately 112° (n = 520 measurements from 5 particles). These results indicate that L137 adopts a flexible three-lobed architecture in solution, consistent with the AF3-predicted monomeric structural model.

**Supplementary Table 1. Data collection and refinement statistics for UDP/Man-bound POMGNT2**

| | Native | $K_2PtCl_4$ |
|---|---|---|
| **Crystallographic data** | | |
| Space group | $P2_12_12_1$ | $P2_12_12_1$ |
| Unit cell *a*/*b*/*c* (Å) | 145.0/150.0/191.5 | 145.4/151.2/191.7 |
| **Data processing statistics** | | |
| Beam line | SPring-8 BL44XU | SPring-8 BL44XU |
| Wavelength (Å) | 0.9000 | 1.0717 |
| Resolution (Å) | 50-2.80 (2.97–2.80) | 48.79-3.71(3.84–3.71) |
| Total/unique reflections | 705,396/197,873 | 1,214,082/45,325 |
| Completeness (%) | 99.7 (99.3) | 99.6 (96.3) |
| $R_{merge}$ (%) | 6.2 (91.8) | 24.8 (184.6) |
| $CC_{1/2}$ | 99.9 (59.6) | 99.9 (75.9) |
| $I / \sigma (I)$ | 13.5 (1.2) | 12.6 (1.8) |
| Multiplicity | 3.6 (3.4) | 26.8 (18.2) |
| **Refinement statistics** | | |
| Resolution (Å) | 47.92-2.80 | |
| $R_{work} / R_{free}$ (%) | 19.1/22.8 | |
| R.m.s. deviations from ideal | | |
| Bond lengths (Å) | 0.008 | |
| Bond angles (°) | 1.63 | |
| Ramachandran plot (%) | | |
| Favored | 93.9 | |
| Allowed | 5.5 | |
| Disallowed | 0.6 | |
| Number of atoms | | |
| Protein atoms (A/B/C/D) | 4192/4121/4163/4157 | |
| UDP | 25/25/25/25 | |
| Mannose | 12/12/12/12 | |

| | |
|---|---|
| Average *B* factors ($Å^2$) | |
| Protein atoms | 88.1/90.7/95.2/102.3 |
| UDP | 104.7/153.5/133.7/147.5 |
| Mannose | 97.7/111.5/108.5/116.6 |
